# A method for detection of permeation events in Molecular Dynamics simulations of lipid bilayers

**DOI:** 10.1101/2021.01.20.427278

**Authors:** Carlos R. S. Camilo, J. Roberto Ruggiero, Alexandre S. de Araujo

## Abstract

The cell membrane is one of the most important structures of life. Understanding its functioning is essential for several human knowledge areas, mainly how it controls the efflux of substances between the cytoplasm and the environment. Being a complex structure, composed of several classes of compounds such as lipids, proteins, sugars, etc., a convenient way to mimic it is through a phospholipid bilayer. The Molecular Dynamics simulation of lipid bilayers in solution is the main computational approach to model the cell membrane. In this work, we present a method to detect permeation events of molecules through the lipid bilayer, characterizing its crossing time and trajectory. By splitting the simulation box into well-defined regions, the method distinguishes the passage of molecules through the bilayer from artifacts produced by crossing molecules through the simulation box edges when using periodic boundary conditions. We apply the method to study the spontaneous permeation of water molecules through bilayers with different lipid compositions and modeled with different force fields. Our method successfully characterizes the permeation events, and the results obtained show that the frequency and time of permeation are independent of the force field used to model the phospholipids. Besides, it is observed that the increase in the concentration of cholesterol molecules in lipid bilayers induces the reduction of permeation events due to its compacting action on the bilayer, making it denser and, therefore, hindering the diffusion of water molecules inside it. The computational tool to perform the method discussed here is available on https://github.com/crobertocamilo/MD-permeation.

## INTRODUCTION

Computational simulations of biological systems by Molecular Dynamics (MD) provide the time evolution at atomic or molecular resolution levels. Analysis of the system and its components can contribute with information about the interrelationship between them and the mechanisms of molecular action.^1^ Bilayers simulations mimic the basic features of biological membranes and have been used to obtain valuable information on lipids interactions and structure. From the trajectories, it is possible to understand how the lipid composition and proportion,^2–4^ as soon as their interaction with drugs, small ions, or other biological interest molecules,^5–7^ affects the bilayer structural properties. Particularly noteworthy are the interactions of water molecules with lipids because they induce the formation and stabilize the bilayer structure.^8,9^

Biological membranes are structures that compartmentalize environments in the cells separating the cytosol from the extracellular environment. In organelles, they provide a distinct biochemical domain from that in the cellular environment.^10^ Although its primary structural function is to act as a barrier between interior and exterior, the cellular activity demands that the membranes are selectively permeable, presenting exchange mechanisms for molecules between the cytosol and the extracellular environment, usually through processes mediated by proteins integrated to the membrane structure. However, the passage of small molecules through lipid membranes is verified experimentally^11^ and in computational simulations,^12,13^ even in the absence of proteins. The two main mechanisms proposed to explain the permeation of ions or small molecules through membranes are the passage through a transient pore established by local disturbances such as solvated ions or the passage directly through the membrane by diffusive processes as observed for H_2_O, O_2_, CO_2_ molecules.^14^ The non-facilitated permeation of small molecules is associated with their solubility in the aqueous phase. If the molecule does not establish strong interactions with water, there is a significant probability that its partition coefficient favors the interaction with the membrane. Once inside it, interaction with lipids may eventually result in the permeation through membrane.^15^ The structure of a membrane presents, in addition to the hydrophobic nucleus defined by the acyl chains, two hydrated interface regions defined by the polar lipid headgroups, which can induce the non-facilitated permeation also of small hydrophilic molecules.

Frequently, to obtain characteristic properties for diffusive phenomena by simulations, such as the permeability coefficient of the interesting molecule, requires enhanced sampling techniques, such as umbrella sampling,^16^ adaptive biasing force (ABF),^17^ or metadynamics.^18,19^ Nevertheless, even in equilibrium MD simulation of a lipid bilayer in the aqueous phase, water molecule permeation events are also observed.^15,20^ The characterization of such events, mainly their frequency of occurrence and duration, can provide relevant information about how the lipid composition of the bilayer affects the permeation of a solute, or if the interaction of a target molecule with the membrane, like a drug, can favor the extravasation of water and other molecules. However, the number of water molecules in the simulated system (typically several dozens for each lipid molecule) and the number of simulation frames make it impossible to check and study permeation events using conventional analysis such as density profiles. Besides, even when it is possible to check the presence of water molecules inside the bilayer, density profiles are time averages and are not suitable to find whether these molecules have completed the passage through the structure, or they have returned to the water bulk through the same leaflet they had entered.

The identification of permeation events requires the analysis of the time evolution of each water molecule position relative to the bilayer, frame by frame, and the application of an algorithm that calculates whether the molecules that entered the bilayer completed or not the passage through it. Furthermore, statistically, any water molecule can enter the lipid structure at some point at the simulation.

In this work, we present a simple method to analyze bilayer simulations trajectories to detect permeation events by solutes or water molecules. Its application to the simulation of a bilayer in aqueous phase, for instance, reveals how many permeation events occurred, which molecules were involved, and their time map. The method can also be applied to bilayer containing integrated proteins or other molecules, providing information about their influence on bilayer permeation during the simulation, which is important to verify hydrated pore formation. To demonstrate the applicability of the method, we present and compare the results of pure POPC (*1-palmitoyl-2-oleoyl-sn-glycero-3-phosphocholine*), POPE (*1-palmitoyl-2-oleoyl-sn-glycero-3-phosphoethanolamine*) and POPG (*1-palmitoyl-2-oleoyl-sn-glycero-3-phosphoglycerol*) bilayer simulations produced in two different MD simulations packages, using two different force fields and distinct molecular models for water. The choice of the two sets has a secondary objective of comparing the effects of different water molecules representation on the water permeation events. Applied to simulations of mixed POPC/Cholesterol (Chol.) bilayers at different concentrations (POPC:Chol. 100:0, 80:20 and 60:40), the method reveals a gradual reduction in the water molecule permeation with increasing cholesterol concentration.

## II. THEORY AND METHODS

### II.A. COMPUTATIONAL DETAILS

The application of the method presented in the next sections requires a molecular dynamics trajectory of the system under study. We use two different sets of MD packages and force fields to generate trajectories, as described below.

The first set of simulations was produced with the GROMACS package,^21,22^ using the united atoms GROMO96 53a6 force field,^23^ modified with the Berger parameters^24^ for lipid molecules representation, denoted as GG. The structure and topology used for the POPC and POPE molecules are distributed by Tieleman;^25^ for POPG they are created by Zhao et al.,^26^ while for the cholesterol molecules, they are the same used by Höltje et al.^27^ The SPC model^28^ was used for the water molecules representation. In the POPG bilayer simulation formed by anionic molecules, Na+ counterions were added to ensure the neutrality of the system. The standard parameters of the force field were used for both water and Na^+^. The rectangular simulation boxes, around 60×60×85 Å, containing a bilayer with 130 lipids (65 per monolayer) and approximately 4600 water molecules (about 35 per lipid molecule) were built using the facilities of the GROMACS package. All the simulation boxes were submitted to a 10 ns period of equilibration and then to a 300 ns period of production dynamics, performed with an integration step of 2 fs, and the trajectories being recorded every 10 ps.

Three-dimensional periodic boundary conditions were applied to all the simulation boxes. Non bounded interactions were treated using the PME algorithm^29^ for long-range electrostatic interactions and a 12 Å cutoff radius for short-range interactions. The LINCS algorithm^30^ was used to control the lengths of atomic bonds in molecules, and the SETTLE algorithm^31^ used to preserve the geometry of water molecules. The simulations were developed in the NPT ensemble, with a reference temperature of 310K, controlled through the Nosé-Hoover thermostat,^32,33^ with coupling for two independent groups (water/ions and lipids), and a reference pressure of 1 bar (0.987 atm) ensured by the Parrinello-Rahman barostat,^34,35^ with semi-isotropic coupling (with the bilayer plane *xy* coupled, but independent *z*). The reference value for the system temperature is above the phase transition temperature of all studied bilayers,^36,37^ ensuring that the analyzed events occur with the bilayers in the liquid-crystalline phase.

The second set of simulations was carried out with the NAMD software package^38^ using the all-atoms CHARMM36 force field,^39^ termed NC. The rectangular simulation boxes, with about the same size of the other set, containing the initial structure of the simulation were built using the CHARMM-GUI Membrane Builder module,^40^ following the same pattern as in the first set i.e., bilayers with 130 lipids and approximately 4600 water molecules, here represented by the TIP3P water model.^41,42^ These simulations also ran for a 10 ns equilibration period followed by a production dynamics of 300 ns. The timestep of 2 fs was used to integrate the dynamical equations, recording the trajectories each 10 ps. Periodic boundary conditions and the non-bonded interaction were treated exactly as in the first set of simulations. The reference temperature and pressure, 310 K and 1 bar, were controlled using Langevin dynamics and the Nosé-Hoover Langevin barostat,^43,44^ respectively. The RATTLE algorithm^45^ was used to restrict covalent bonds with hydrogen atoms and the SETTLE algorithm to preserve the geometry of water molecules.

### II.B. PERMEATION: A METHOD FOR EVENT ANALYSIS

In this work, we present a simple application method for identifying permeation events of molecules (e.g. H_2_O or O_2_) in an MD bilayer simulation. The method consists of analyzing the position of the solute molecule relative to the bilayer and characterizing permeation events. Its application to all solute or solvent molecules provides the mapping of all permeation events that occur during a simulation. Here it will be applied for water molecules.

Before we describe the method, let us follow the *z* coordinate of a chosen water molecule throughout the simulation of a phospholipid bilayer of POPC in an aqueous medium, shown in Figure 1a. The water molecule diffuses on the aqueous phase and eventually enters the hydrated phase of the bilayer, shown as the orange lines, and defined by the averaged *z* coordinate of the phosphate groups center of mass at each instant of the simulation. Note that dozens of occurrences are observed. They vary in duration and depth, i.e., how long the molecule remains inside the structure and how much the molecule enters the lipid phase towards the bilayer center (dashed line in yellow). Around the instant 280 ns, one particular event draws attention: the molecule enters the bilayer from the lower interface and advances through the center of the structure and, after a few ns, leaves the bilayer by the top interface. Details of this interval using a line representation for the movement between two registered instants are shown in Figure 1b, where the trajectory of the tracked water molecule reveals the specific position and time of its passage through the lipid structure. In an analogy with the biological system, the molecule would have gone from the cytosol to the extracellular environment, or vice versa, crossing the cell membrane. Therefore, we have documented a permeation event of a water molecule through a bilayer, in a process that occurs by non-facilitated diffusion, i.e., during the dynamic of an equilibrium simulation, without including any explicit inducing potential or gradient of chemical potential.

**Figure 1.**
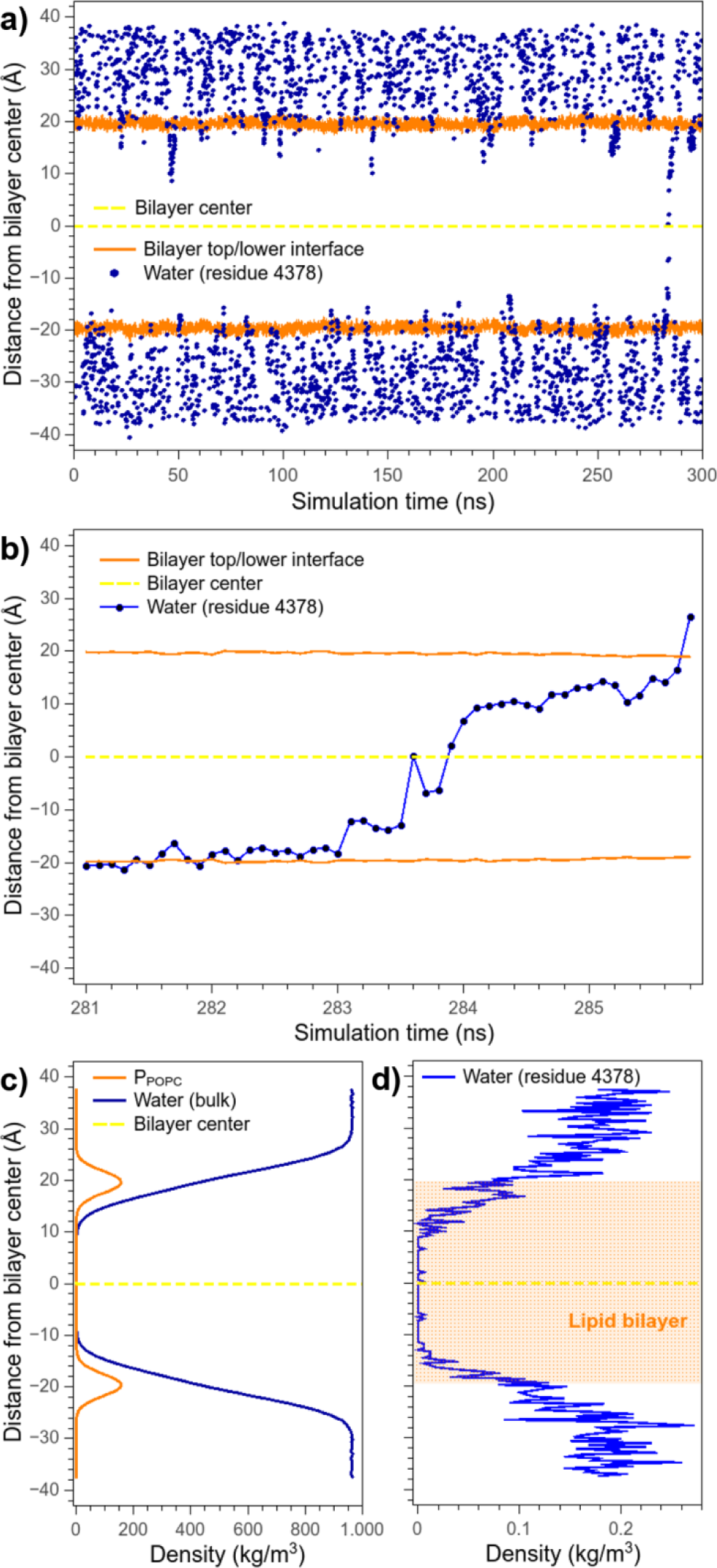
(a) Temporal evolution of the *z*-coordinate of a single water molecule (blue dots) in a simulation box containing a POPC bilayer in the aqueous phase. The upper and lower limits of the bilayer (orange lines) are the average *z* coordinates of the phospholipids P atoms at each instant of the simulation. The geometric center of the lipidic structure is shown as dashed yellow lines. (b) A highlight of a water molecule permeation event occurring at 281 to 286 ns time interval of (a) (here a line connects the dots. (c) Density profile for the P atoms of the POPC molecules and all water molecules in the simulation box, normalized relative to the bilayer center. There is no evidence of water inside the hydrophobic nucleus of the bilayer. (d) Individual density profile of the water molecule residue analyzed in (a) and (b) – the tiny peaks in the central region reveals that the molecule was inside the bilayer, but the information is not enough to characterize the event highlighted in (b).

The usual way of analyzing the distribution of a chemical group or molecule across the simulation box is by calculating its density profile along the axis perpendicular to the bilayer (typically, the *z*-axis).^9,26,46,47^ From Figure 1c, however, no permeation event can be identified from the water density profile. Even when applied to the specific water residue (Figure 1d), which we have already identified to perform a permeation event, the obtained distribution provides only a tiny indication that the molecule was inside the bilayer during the analyzed interval. Nevertheless, it is not possible to follow the temporal evolution of the phenomenon. Questions such as whether the molecule indeed passes through the structure, or advances to the center but returns, cannot be answered using density profiles analysis. Therefore, even if it was applied individually to each water molecule in the simulation box, density profiles cannot detect permeation events. If an event consists of just a few points in the trajectory (short time duration), information about their occurrence can hardly be obtained by visual inspection of density profile graphics or using software to visualize the trajectory of the molecules. In this way, it is evident that such events require an individual analysis for each molecule of interest, so it is convenient to use some specific analysis method for that purpose.

The structure of a bilayer defines, along the axis perpendicular to its plane (z-axis), regions with significantly different physical properties (see Figure 2a):

**Figure 2.**
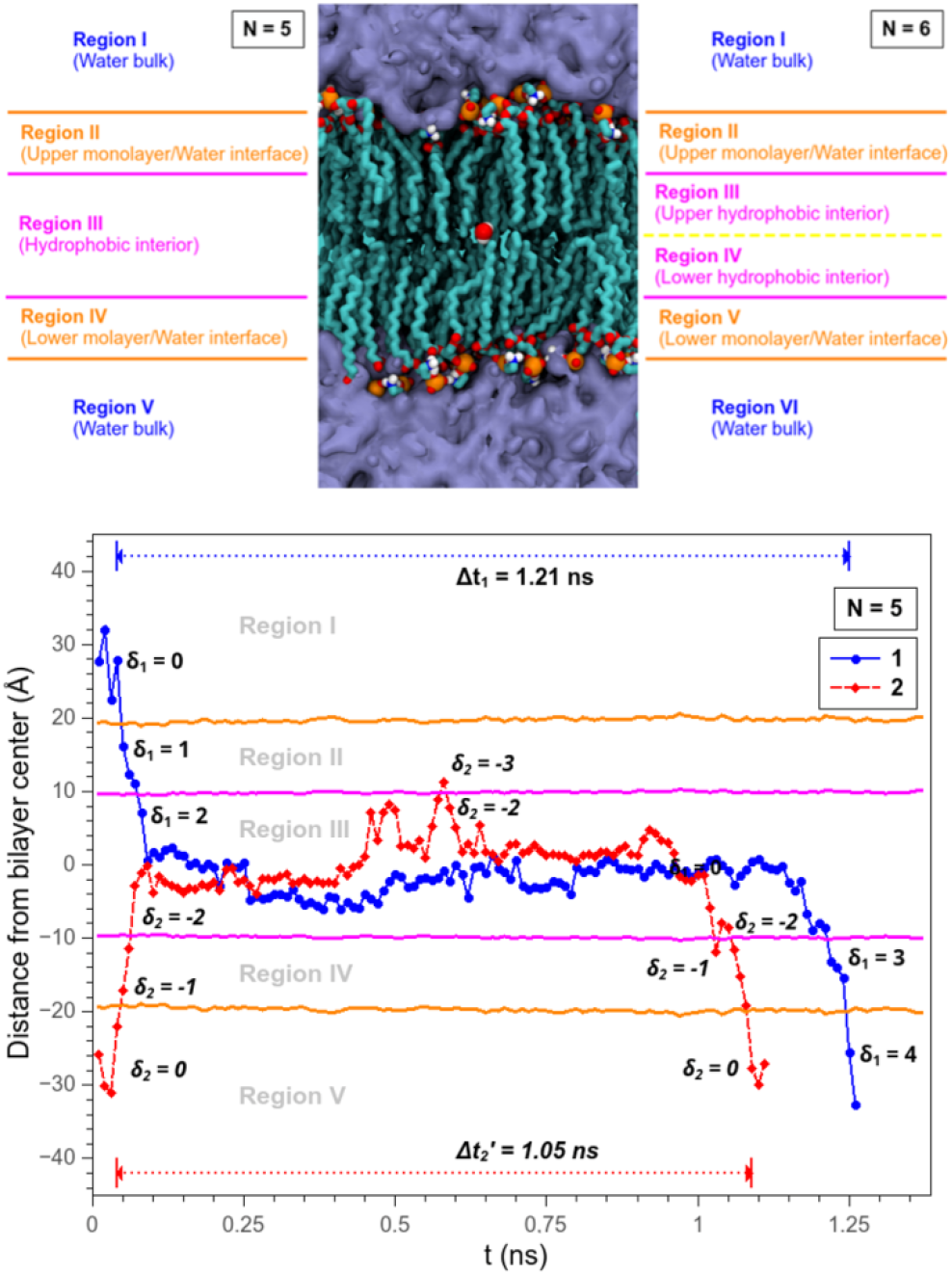
(a) Example of segmentation of a simulation box in five and six regions. The blue bulk represents the aqueous phase, the orange spheres are P atoms of the phospholipid headgroups and the apolar tails are presented in cyan. An individual water molecule, located in the bilayer center, is highlighted in white (hydrogens) and red (oxygen). (b) Examples of a permeation event (in blue) successfully completed, and another one in which, although the molecule advances over the center of the bilayer, it ends up returning and leaving the structure for the same monolayer it had entered (in red), not completing a passage so. The indication of regions follows the *N* = 5 pattern defined in (a), and the accumulated score value δ of each molecule is highlighted when their region change.

- Region I is the upper aqueous phase, where other polar molecules may also be dissolved.
- Region II is the solvated region of the bilayer, comprising the phospholipids headgroups and glycerol groups.
- Region III comprises: the acyl tails of phospholipids, a highly hydrophobic region where the presence of polar molecules is ordinarily negligible, the center of the bilayer, and the acyl tails region of the opposite monolayer.
- Region IV and V are the same as regions II and I, respectively, at the opposite monolayer.

Different authors segment the bilayer in regions according to their objectives, and systems studied.^15,48^ In any case, a permeation event consists of a molecule gradually advancing through all the bilayer regions. To complete a passage, the molecule must enter by one monolayer and exit on the other. In this work, movements in which the molecule enter and exit through the same monolayer, different from other reference,^20^ are not considered a permeation event. Here we divide the bilayer into the five regions described above, but the method presented below can be generalized for an arbitrary number of regions.

A way to analyze permeation events is to segment the bilayer into different regions in *z* and follow the temporal evolution of each solute molecule in these regions. Using the segmentation in five regions shown in Figure 2a, for instance, when going from Region I to Region II (transition I→II), the molecule will have left the aqueous phase and entered the upper interface region of the bilayer. If it goes further into the structure, the molecule will reach the hydrophobic interior (Region III), but if it returns to the water bulk, it will be again in Region I, without having completed a passage. As seen earlier, events in which the molecule enters and remains in the bilayer for some simulation time, but then leaves the structure by the same interface it had entered are frequent (Figure 1a).

To follow the evolution of the *i*-th molecule trajectory relative to the bilayer, we can define a quantity T_i_(t) that measures the number of region borders transpassed by the particle between two consecutive instants, t and t + Δt. Let Rt. Let R_i_(t) be a label defining the region where the particle is located at the instant t. If in the instant t + Δt the particle remains in the same region, T_i_(t) should be zero, and different from zero in other cases.

Let us introduce, for a five regions segmentation, as shown in Figure 2a, the transition matrix *M*(R_i_(t + Δt),R_i_(t)) defined by

The element of *M* corresponding to the movement between t and t + Δt. Let Rt will give the value of T_i_(t). Note that this matrix permits to follow the motion direction because positive values correspond to transition from upper to lower monolayer while the negatives for movements in the opposite direction.

Due to the periodic boundary conditions applied to MD simulations, Regions I and V constitute the water bulk, and the molecules transact from one to another through the simulation box images. So considering them as distinct regions is just an artifact of the method, and therefore transitions like I→V or V→I must be disregarded. Transitions with values 2 and 3 (T = ± 2 or T = ± 3) imply that from one analyzed instant to another, the molecule *jumps* one or two entire regions, respectively, crossing two or three borders. For example, the molecule was in the upper aqueous phase, and in the next instant, it is in the center of the bilayer, performing a I→III transition. This type of transition is directly associated with the accuracy of the analyzed trajectory (see Figure S1 in the Supporting Information). For water molecules, trajectories recorded at least each 25 ps timeframe were necessary to eliminate the types 2 or 3 *jumps* occurrences. For lower precision recorded trajectories or a higher segmentation of the bilayer, type 2 or greater *jumps* can occur, and the proper permeation event characterization requires an individual graphical analysis, as exemplified in Figure S1.

In a typical permeation event, the molecule progressively advances from one region to another until the passage is complete. Therefore, in addition to monitoring the evolution of the molecule position between consecutive instants, it is necessary to verify when a set of transitions characterizes a permeation event. Considering a segmented box in *N* regions, we will have a permeation event when

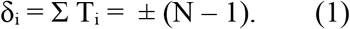

For *N* = 5, a permeation event occurs when a score of 4 is reached (the molecule left Region I and reached Region V by crossing the bilayer) or −4 (starts at Region V and reach Region I). Figure 2b shows two situations, one where a permeation event occurs (blue line) and another in which, despite accessing the center of the bilayer, the molecule returns to the bulk water by the same interface, not completing a permeation event (red line). Once the molecule score has reached the criterion defined above, the event must be registered and imposed δ = 0, since the molecule having completed the passage is again in the aqueous phase.

Some care must be taken at the beginning of an analysis. For all molecules located at Regions I or V, δ_i_ is set 0. The δ value of molecules initially in Regions II to IV is ignored until the molecules reach Regions I or V, unless one provides this information from a previous analysis.

The number *N* of regions depends on the system being analyzed, mainly considering their physical meaning and the studied interest molecule. To evaluate water permeation events in pure bilayers, *N* = 5 is adequate, as shown in the following applications. On the other hand, if one wants to know how many water molecules reach a given region of the bilayer, its center for example, without being a permeation event, *N* = 6 can be adopted considering the bilayer center as a new region boundary (Figure 2a). In this case, for δ = ± 5 we would have the permeation events completed. Denoting Δ the total number of permeation events, and Δ’. the number of events for which water molecules reach the center of the bilayer for the first time (δ reaches ± 3) during its journey inside the bilayer. Then Δ’ - Δ will provide the number of events in which the permeation is incomplete, that is, events where the molecule reached the center, returned, and left the bilayer by the same interface. However, it is essential to highlight the compromise between the number of regions *N* and the recorded trajectory timeframe. Increasing *N* means a decrease in the thickness of the layers defining the different regions, and the frequency of occurrence of *jumps* requiring special treatment will grow. Therefore, increases in *N* need to be compensated by decreasing the timeframe of the analyzed trajectory.

Phospholipid bilayers may show local fluctuations in thickness for several reasons. For example, the interactions with a peptide, or other molecules, can produce a local thinning of the bilayer.^6,49^ A bilayer composed of different types of lipids can also produce fluctuations in the bilayer thickness due to the size of the tails or the number of unsaturations.^50^ Although the definition of regions is made from averages on the position of specific groups (phosphate, carbonyl, etc.), a permeation event implies that the molecule enters by an interface, passes through the structure, and exits by the other interface. In this way, the proposed method can also be applied to bilayers with local deformations analyzing the time evolution of *z* coordinate as a function of *x* and *y*. So, correlating these points and the number of events, one can indicate if the deformation facilitates or promotes the increase of permeation events.

## III. RESULTS AND DISCUSSION

### III.A. PERMEATION OF WATER MOLECULES IN PURE AND MIXED BILAYERS

Biological membranes are composed of several types of lipids and other molecules forming a system whose size makes it very complicated to study directly through computer simulations.^51,52^ On the other hand, phospholipid bilayers are systems that have structural properties of cell membranes, but on an adequate scale to a computational approach.^53^ A pure bilayer is a model membrane composed of a single type of phospholipid. This simplified system is normally used as the starting point for tests or validations. Simplifying the system is also a strategy used in experimental techniques with films or micelles formed by a single lipid type.^54^ Once successfully applied in the pure system, the bilayer composition can be progressively changed to a system formed by more types of lipids, or by the insertion of other molecules of interest. This gradual increase in complexity leads to a more realistic representation of the studied system.^55^

In MD simulations, the system molecules are modeled according to the parameters of a force field. The GROMOS96-Berger and CHARMM36 force fields provide representations that have been widely used in bilayer simulations^56^ (see Computational Details). To demonstrate the applicability of the presented method, we performed simulations of pure POPC, POPE, and POPG bilayers, some of the main phospholipids components of membranes in eukaryotic and prokaryotic cells.^10^

The application of the method to the simulations (see Figure 3) reveals the occurrence of permeation events in all studied bilayers and that they are distributed throughout the simulation time. The red curves show the cumulative histogram of permeation events for the GG simulations, while the black curves are for the NC set. A good correlation is observed in the results obtained in the two sets of simulations for the three pure bilayers. For example, in the POPC bilayers (Figure 3a), a permeation event occurs, on average, every 43 ns of simulation (1/43 ns) for GG and 50 ns (1/50 ns) for NC. In the POPE bilayers (Figure 3b), the average frequencies are 1/75 ns in GG and 1/50 ns for NC. And, in the POPG anionic bilayers (Figure 3c), the simulations show equivalent results but with a smaller number of events, 1/100 ns for GG and 1/75 ns for NC. These results confirm the expected equivalence between the two different sets of simulations, and that the proposed method works properly for both of them.

**Figure 3.**
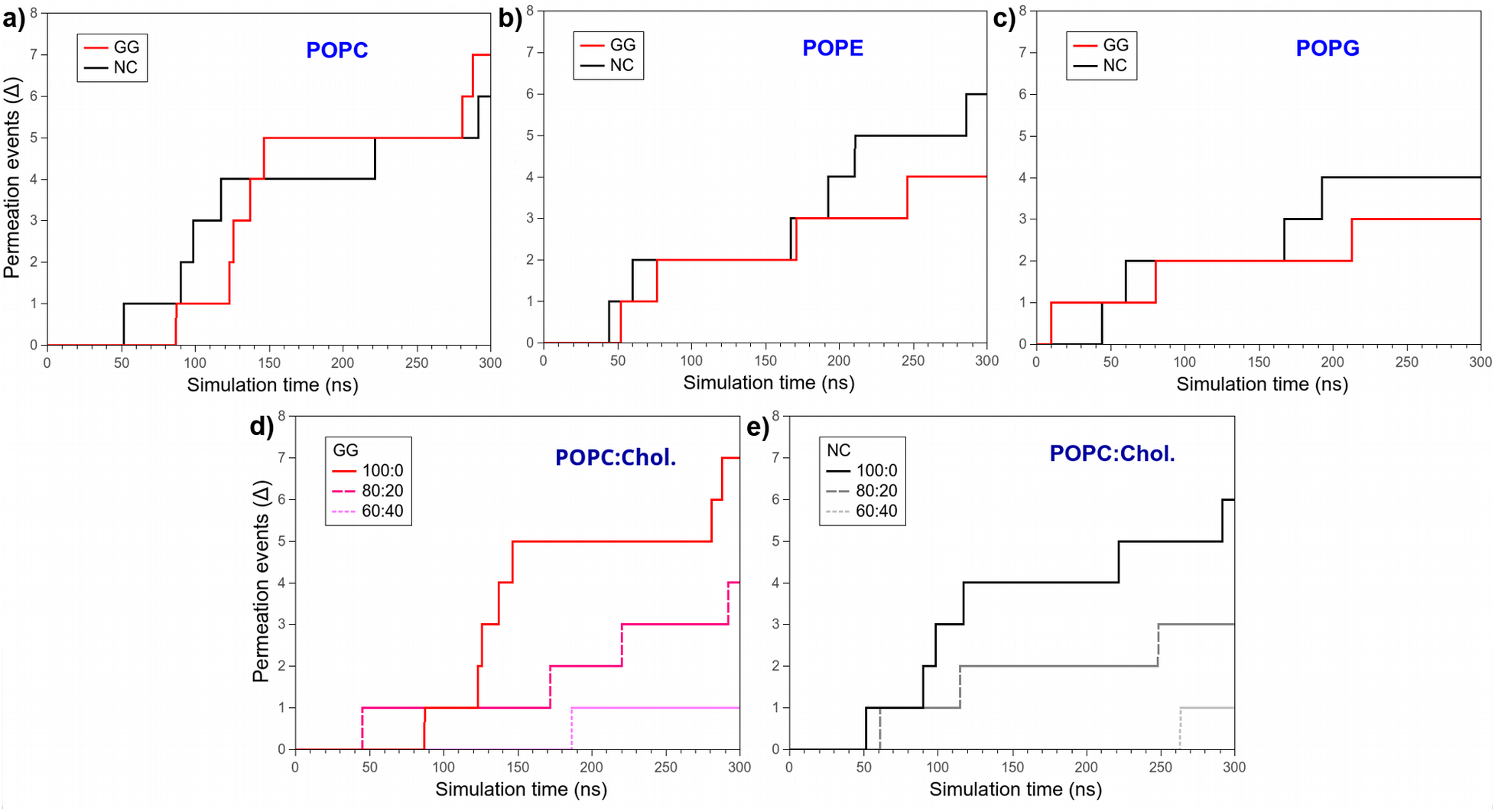
Histogram of the cumulative number of permeation events in pure POPC (a), POPE (b), and POPG (c) bilayers, simulated with different force fields and simulation packages: GROMACS/GROMOS96 (lines in red) and NAMD/CHARMM36 (black lines). Comparison of the number of permeation events in bilayers with different concentrations of cholesterol, POPC:Chol. 100:0 (darkest line), 80:20 and 60:40 (lightest line) obtained with GROMACS/GROMOS96 (d) and NAMD/CHARMM36 (e).

The next step was evaluating the method at binary systems containing two types of lipids. Eukaryotic membranes are cholesterol rich, and its presence brings major modifications to the membrane properties, mainly reducing fluidity and increasing the lipid package (smaller area per lipid) and the thickness of the hydrophobic phase.^57,58^ Starting from a pure system (POPC: Chol. 100: 0) and gradually increasing the proportion of Chol. molecules (80:20 and 60:40) it is clear a proportional reduction in the number of water molecule permeation events. This effect is observed for both sets of simulations (Figures 3d for GG and 3e for NC) with a good agreement between them. While for GG, the average frequency of a permeation event drops from 1/43 ns for pure bilayer (100:0) to 1/75 ns in the bilayer containing 20% Chol. (80:20) and 1/300 ns in 40% Chol. (60:40), for the NC simulations, these numbers are 1/50 ns (100: 0), 1/100 ns (80:40), and 1/300 ns (60:40).

The results indicate that Chol. has an inhibitory effect on water permeation. They point the same qualitative findings of Hub et al^59^ that, using statistical sampling simulations, show an increase in the transfer free energy barrier of water molecules through a bilayer containing cholesterol molecules. Also, Jedlovszky and Mezei^60^ show a shift outwards the membrane due to the enlargement of the free energy barrier with the increasing of cholesterol content. These changes in the bilayer thermodynamics reduce the water solubility at the hydrophobic interior, thicker and denser, and consequently decrease the probability of a water molecule reaches the center of the structure.

Using a different condition to evaluate the accessibility of water molecules to the bilayer interior in microsecond-lengthscale equilibrium simulations, Hong et al.^20^ also report a reduction in the number of events with Chol. insertion. Using the VMD,^61^ the authors considered as permeation event every water molecule that could be located at some instant in a central region of ± 9 Å from the bilayer center, without following its temporal evolution. While for a pure POPC bilayer they obtained one event every 17 ns, in a system containing 33% Chol. the average dropped to 1/114 ns, that is, a 6.7x reduction in the frequency of events, which can be located between the reduction we obtained to 20% Chol. (1.7x GG or 2.0x NC) and 40% Chol. (7.0x GG and NC). However, some care must be taken to establish direct comparisons between the number of permeation events obtained here with those reported by the authors, since the criteria used to define an event are different. Imposing a less restrictive condition to compute permeation events, that the water molecules reach the center of the bilayer, measured as Δ’ in the data of Table 2, we still obtained about the same factors for reduction of the frequencies pointed above. Interestingly, the results reveal that around half of the water molecules cross the bilayer center and half arrive until the center but return to the bulk by the same monolayer. In Table 2, Δ and ΔT measure the number and the average duration of complete permeation events, respectively, while Δ’ and ΔT’ measure the same quantities for all the events in which the water molecule reaches the bilayer center. The data indicate that once in the bilayer center, the probability of the molecule completing the crossing by the opposite monolayer, or returning through the same one it had entered, is statistically equivalent. This result is consistent with the free energy profiles obtained through enhanced sampling simulations for pure bilayers with small curvatures, which show that the center of the structure as a local minimum between two symmetrical peaks.^62,63^

**Table 1.** Transition values for *N* = 5 regions.

|  |  | R (t) |  |  |  |  |
| --- | --- | --- | --- | --- | --- | --- |
|  |  | I | II | III | IV | V |
| R (t + $\Delta t$) | I | 0 | 1 | 2 | 3 | 0 |
|  | II | -1 | 0 | 1 | 2 | 3 |
|  | III | -2 | -1 | 0 | 1 | 2 |
|  | IV | -3 | -2 | -1 | 0 | 1 |
|  | V | 0 | -3 | -2 | -1 | 0 |

**Table 2.**
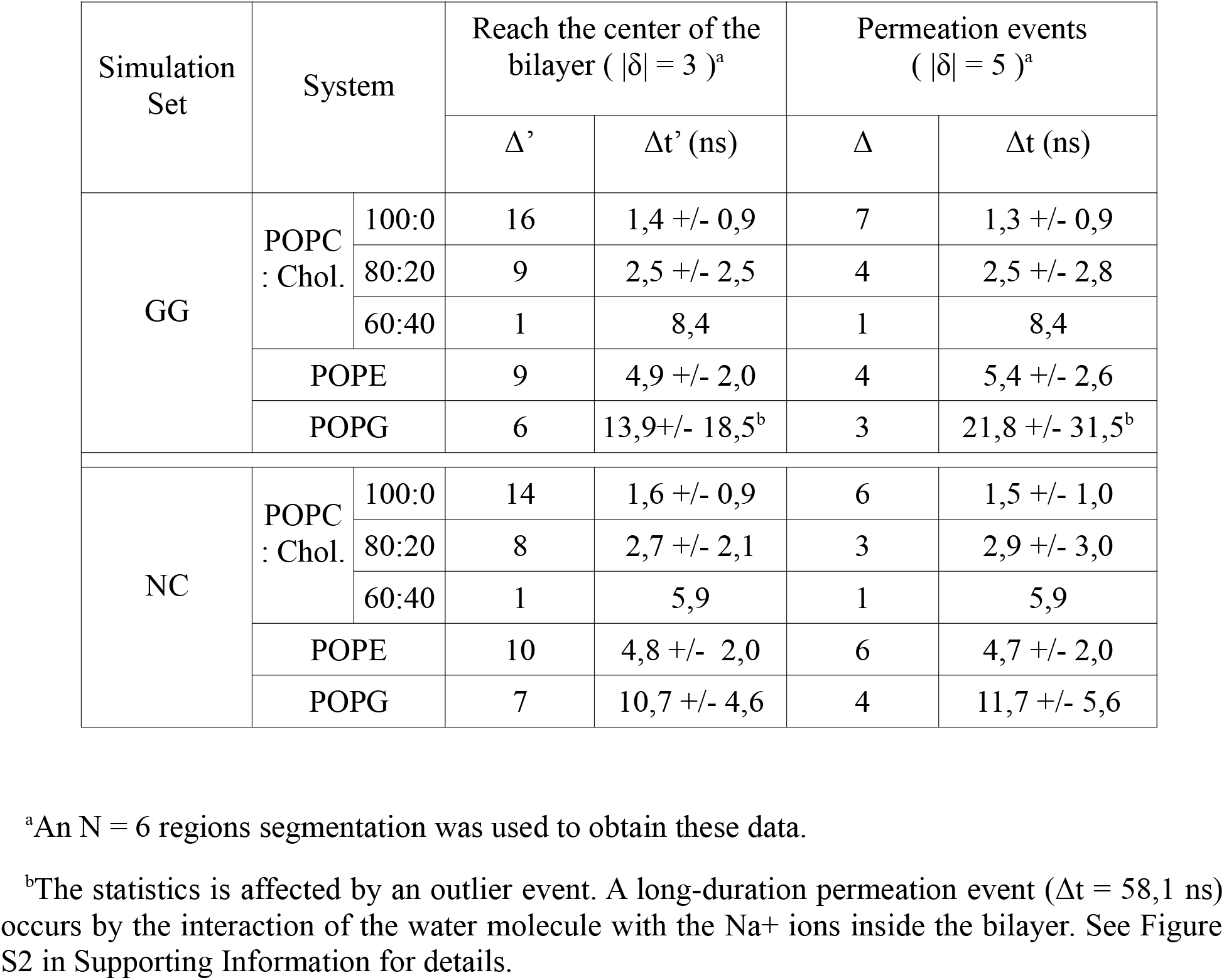
Comparison between the number of molecules that reach the center of the bilayer (Δ’) with those between them that complete permeation events (Δ). The time intervals are the total simulation time the molecule remains within the structure.

The results show that the number of permeation events decreases, for pure bilayers, following the same trend as the area per lipid, i.e., POPC→POPE→POPG. The same happens to the bilayers with cholesterol. For more reliable results, longer simulation time should be used to improve the statistical sampling. However, simulations of a few hundred nanoseconds can provide a qualitative insight about the effect of a certain molecule on the bilayer, as cholesterol, and, consequently, on the number and duration of permeation events. Finally, as expected for equilibrium MD simulations, we observed an approximately equal number of permeation events in both directions.

## IV. CONCLUSION

The passage of small molecules through membranes by diffusive processes is a phenomenon that can also be studied by MD simulations. Even in simulations without a chemical potential gradient or osmotic pressure, permeation events can occur. In this work, a simple method to identify the occurrence of molecule permeation events in lipid bilayers was presented. The method provided the mapping of all permeation events during the simulation, when applied to MD trajectories for lipid bilayers in aqueous solutions.

To illustrate the method, we present water permeation investigation in pure bilayer simulations carried out using different force fields and simulation packages. The results obtained indicate an equivalence between these setups. Applied to the mixed POPC:Chol. bilayers, the method shows that the increase in cholesterol concentration gradually reduces the occurrence of water molecule permeation events. From these data, it is possible to make statistical analyzes of the frequency of event occurrence and their duration intervals, as shown.

In bilayers disturbed by the action of a drug or membrane protein, for example, the distribution analysis of events throughout the simulation time can provide clues about the interaction dynamics between bilayer and drug. By analyzing the x, y coordinates of the water molecules during permeations, it is possible to evaluate whether and how the disturbances favor the extravasation of water.

## Supporting information

Supplementary Graphics

## ASSOCIATED CONTENT

### Supporting Information

The following files are available free of charge.

Influence of the trajectory precision over the detection of permeation events by examples in which occur *jumps*.

## AUTHOR INFORMATION

### Author Contributions

CRC performed all simulations and analysis. The manuscript was written through contributions of all authors. All authors have given approval to the final version of the manuscript.

## Notes

The authors declare no competing financial interest.

## ACKNOWLEDGMENT

We would like to thank to Pedro G. Pascutti for the motivation.

## References

(1) Gumbart, J., Aksimentiev, A., Tajkhorshid, E., Wang, Y., Schulten, K. Molecular Dynamics Simulations of Proteins in Lipid Bilayers. Curr Opin Struct Biol 2005, 15 (4), 423– 431. https://doi.org/10.1016/j.sbi.2005.07.007.

(2) Tieleman, D. P., Marrink, S. J., Berendsen, H. J. A Computer Perspective of Membranes: Molecular Dynamics Studies of Lipid Bilayer Systems. Biochim. Biophys. Acta 1997, 1331 (3), 235–270. https://doi.org/10.1016/s0304-4157(97)00008-7.

(3) Marrink, S. J., de Vries, A. H., Tieleman, D. P. Lipids on the Move: Simulations of Membrane Pores, Domains, Stalks and Curves. Biochimica et Biophysica Acta (BBA) – Biomembranes 2009, 1788 (1), 149–168. https://doi.org/10.1016/j.bbamem.2008.10.006.

(4) Moradi, S., Nowroozi, A., Shahlaei, M. Shedding Light on the Structural Properties of Lipid Bilayers Using Molecular Dynamics Simulation: A Review Study. RSC Adv. 2019, 9 (8), 4644–4658. https://doi.org/10.1039/C8RA08441F.

(5) Deol, S. S., Bond, P. J., Domene, C., Sansom, M. S. P. Lipid-Protein Interactions of Integral Membrane Proteins: A Comparative Simulation Study. Biophysical Journal 2004, 87 (6), 3737–3749. https://doi.org/10.1529/biophysj.104.048397.

(6) Kopeć, W., Telenius, J., Khandelia, H. Molecular Dynamics Simulations of the Interactions of Medicinal Plant Extracts and Drugs with Lipid Bilayer Membranes. The FEBS Journal 2013, 280 (12), 2785–2805. https://doi.org/10.1111/febs.12286.

(7) Ulmschneider, J. P., Ulmschneider, M. B. Molecular Dynamics Simulations Are Redefining Our View of Peptides Interacting with Biological Membranes. Acc. Chem. Res. 2018, 51 (5), 1106–1116. https://doi.org/10.1021/acs.accounts.7b00613.

(8) Nickels, J. D., Katsaras, J. Water and Lipid Bilayers. In Membrane Hydration: The Role of Water in the Structure and Function of Biological Membranes; Disalvo, E. A., Ed., Subcellular Biochemistry; Springer International Publishing: Cham, 2015; pp 45–67. https://doi.org/10.1007/978-3-319-19060-0_3.

(9) Pasenkiewicz-Gierula, M., Baczynski, K., Markiewicz, M., Murzyn, K. Computer Modelling Studies of the Bilayer/Water Interface. Biochimica et Biophysica Acta (BBA) – Biomembranes 2016, 1858 (10), 2305–2321. https://doi.org/10.1016/j.bbamem.2016.01.024.

(10) Alberts, B., Johnson, A., Lewis, J., Raff, M., Roberts, K., Walter, P. The Compartmentalization of Cells. Molecular Biology of the Cell. 4th edition 2002.

(11) Finkelstein, A. Water and Nonelectrolyte Permeability of Lipid Bilayer Membranes. J. Gen. Physiol. 1976, 68 (2), 127–135. https://doi.org/10.1085/jgp.68.2.127.

(12) Orsi, M., Sanderson, W. E., Essex, J. W. Permeability of Small Molecules through a Lipid Bilayer: A Multiscale Simulation Study. J. Phys. Chem. B 2009, 113 (35), 12019–12029. https://doi.org/10.1021/jp903248s.

(13) Venable, R. M., Krämer, A., Pastor, R. W. Molecular Dynamics Simulations of Membrane Permeability. Chem. Rev. 2019, 119 (9), 5954–5997. https://doi.org/10.1021/acs.chemrev.8b00486.

(14) Volkov, A. G., Paula, S., Deamer, D. W. Two Mechanisms of Permeation of Small Neutral Molecules and Hydrated Ions across Phospholipid Bilayers. Bioelectrochemistry and Bioenergetics 1997, 42 (2), 153–160. https://doi.org/10.1016/S0302-4598(96)05097-0.

(15) Awoonor-Williams, E., Rowley, C. N. Molecular Simulation of Nonfacilitated Membrane Permeation. Biochimica et Biophysica Acta (BBA) – Biomembranes 2016, 1858 (7, Part B), 1672–1687. https://doi.org/10.1016/j.bbamem.2015.12.014.

(16) Kästner, J. Umbrella Sampling. WIREs Computational Molecular Science 2011, 1 (6), 932–942. https://doi.org/10.1002/wcms.66.

(17) Darve, E., Pohorille, A. Calculating Free Energies Using Average Force. J. Chem. Phys. 2001, 115 (20), 9169–9183. https://doi.org/10.1063/1.1410978.

(18) Laio, A., Parrinello, M. Escaping Free-Energy Minima. PNAS 2002, 99 (20), 12562– 12566. https://doi.org/10.1073/pnas.202427399.

(19) Valsson, O., Tiwary, P., Parrinello, M. Enhancing Important Fluctuations: Rare Events and Metadynamics from a Conceptual Viewpoint. Annual Review of Physical Chemistry 2016, 67 (1), 159–184. https://doi.org/10.1146/annurev-physchem-040215-112229.

(20) Hong, C., Tieleman, D. P., Wang, Y. Microsecond Molecular Dynamics Simulations of Lipid Mixing. Langmuir 2014, 30 (40), 11993–12001. https://doi.org/10.1021/la502363b.

(21) Berendsen, H. J. C., van der Spoel, D., van Drunen, R. GROMACS: A Message-Passing Parallel Molecular Dynamics Implementation. Computer Physics Communications 1995, 91 (1), 43–56. https://doi.org/10.1016/0010-4655(95)00042-E.

(22) Abraham, M. J., Murtola, T., Schulz, R., Páll, S., Smith, J. C., Hess, B., Lindahl, E. GROMACS: High Performance Molecular Simulations through Multi-Level Parallelism from Laptops to Supercomputers. SoftwareX 2015, 1–2, 19–25. https://doi.org/10.1016/j.softx.2015.06.001.

(23) Oostenbrink, C., Villa, A., Mark, A. E., van Gunsteren, W. F. A Biomolecular Force Field Based on the Free Enthalpy of Hydration and Solvation: The GROMOS Force-Field Parameter Sets 53A5 and 53A6. J Comput Chem 2004, 25 (13), 1656–1676. https://doi.org/10.1002/jcc.20090.

(24) Berger, O., Edholm, O., Jähnig, F. Molecular Dynamics Simulations of a Fluid Bilayer of Dipalmitoylphosphatidylcholine at Full Hydration, Constant Pressure, and Constant Temperature. Biophys. J. 1997, 72 (5), 2002–2013. https://doi.org/10.1016/S0006-3495(97)78845-3.

(25) Tieleman, D. P., Berendsen, H. J. A Molecular Dynamics Study of the Pores Formed by Escherichia Coli OmpF Porin in a Fully Hydrated Palmitoyloleoylphosphatidylcholine Bilayer. Biophys. J. 1998, 74 (6), 2786–2801. https://doi.org/10.1016/S0006-3495(98)77986-X.

(26) Zhao, W., Róg, T., Gurtovenko, A. A., Vattulainen, I., Karttunen, M. Atomic-Scale Structure and Electrostatics of Anionic Palmitoyloleoylphosphatidylglycerol Lipid Bilayers with Na+ Counterions. Biophys. J. 2007, 92 (4), 1114–1124. https://doi.org/10.1529/biophysj.106.086272.

(27) Höltje, M., Förster, T., Brandt, B., Engels, T., von Rybinski, W., Höltje, H. D. Molecular Dynamics Simulations of Stratum Corneum Lipid Models: Fatty Acids and Cholesterol. Biochim. Biophys. Acta 2001, 1511 (1), 156–167. https://doi.org/10.1016/s0005-2736(01)00270-x.

(28) Berendsen, H. J. C., Postma, J. P. M., van Gunsteren, W. F., Hermans, J. Interaction Models for Water in Relation to Protein Hydration. In Intermolecular Forces: Proceedings of the Fourteenth Jerusalem Symposium on Quantum Chemistry and Biochemistry Held in Jerusalem, Israel, April 13–16, 1981; Pullman, B., Ed., The Jerusalem Symposia on Quantum Chemistry and Biochemistry; Springer Netherlands: Dordrecht, 1981; pp 331–342. https://doi.org/10.1007/978-94-015-7658-1_21.

(29) Darden, T., York, D., Pedersen, L. Particle Mesh Ewald: An N·log(N) Method for Ewald Sums in Large Systems. J. Chem. Phys. 1993, 98 (12), 10089–10092. https://doi.org/10.1063/1.464397.

(30) Hess, B., Bekker, H., Berendsen, H. J. C., Fraaije, J. G. E. M. LINCS: A Linear Constraint Solver for Molecular Simulations. Journal of Computational Chemistry 1997, 18 (12), 1463–1472. https://doi.org/10.1002/(SICI)1096-987X(199709)18:12<1463::AID-JCC4>3.0.CO;2-H.

(31) Miyamoto, S., Kollman, P. A. Settle: An Analytical Version of the SHAKE and RATTLE Algorithm for Rigid Water Models. Journal of Computational Chemistry 1992, 13 (8), 952–962. https://doi.org/10.1002/jcc.540130805.

(32) Nosé, S. A Molecular Dynamics Method for Simulations in the Canonical Ensemble. Molecular Physics 1984, 52, 255–268. https://doi.org/10.1080/00268978400101201.

(33) Hoover, W. G. Canonical Dynamics: Equilibrium Phase-Space Distributions. Phys. Rev. A 1985, 31 (3), 1695–1697. https://doi.org/10.1103/PhysRevA.31.1695.

(34) Parrinello, M., Rahman, A. Polymorphic Transitions in Single Crystals: A New Molecular Dynamics Method. Journal of Applied Physics 1981, 52 (12), 7182–7190. https://doi.org/10.1063/1.328693.

(35) Nosé, S., Klein, M. L. Constant Pressure Molecular Dynamics for Molecular Systems. Molecular Physics 1983, 50 (5), 1055–1076. https://doi.org/10.1080/00268978300102851.

(36) Halling, K. K., Ramstedt, B., Nyström, J. H., Slotte, J. P., Nyholm, T. K. M. Cholesterol Interactions with Fluid-Phase Phospholipids: Effect on the Lateral Organization of the Bilayer. Biophysical Journal 2008, 95 (8), 3861–3871. https://doi.org/10.1529/biophysj.108.133744.

(37) Leekumjorn, S., Sum, A. K. Molecular Characterization of Gel and Liquid-Crystalline Structures of Fully Hydrated POPC and POPE Bilayers. J. Phys. Chem. B 2007, 111 (21), 6026–6033. https://doi.org/10.1021/jp0686339.

(38) Klauda, J. B., Venable, R. M., Freites, J. A., O’Connor, J. W., Tobias, D. J., Mondragon-Ramirez, C., Vorobyov, I., MacKerell, A. D., Pastor, R. W. Update of the CHARMM All-Atom Additive Force Field for Lipids: Validation on Six Lipid Types. J. Phys. Chem. B 2010, 114 (23), 7830–7843. https://doi.org/10.1021/jp101759q.

(39) Phillips, J. C., Braun, R., Wang, W., Gumbart, J., Tajkhorshid, E., Villa, E., Chipot, C., Skeel, R. D., Kalé, L., Schulten, K. Scalable Molecular Dynamics with NAMD. J Comput Chem 2005, 26 (16), 1781–1802. https://doi.org/10.1002/jcc.20289.

(40) Jo, S., Lim, J. B., Klauda, J. B., Im, W. CHARMM-GUI Membrane Builder for Mixed Bilayers and Its Application to Yeast Membranes. Biophys. J. 2009, 97 (1), 50–58. https://doi.org/10.1016/j.bpj.2009.04.013.

(41) Jorgensen, W. L., Chandrasekhar, J., Madura, J. D., Impey, R. W., Klein, M. L. Comparison of Simple Potential Functions for Simulating Liquid Water. J. Chem. Phys. 1983, 79 (2), 926–935. https://doi.org/10.1063/1.445869.

(42) Durell, S. R., Brooks, B. R., Ben-Naim, A. Solvent-Induced Forces between Two Hydrophilic Groups. J. Phys. Chem. 1994, 98 (8), 2198–2202. https://doi.org/10.1021/j100059a038.

(43) Martyna, G. J., Tobias, D. J., Klein, M. L. Constant Pressure Molecular Dynamics Algorithms. J. Chem. Phys. 1994, 101 (5), 4177–4189. https://doi.org/10.1063/1.467468.

(44) Feller, S. E., Zhang, Y., Pastor, R. W., Brooks, B. R. Constant Pressure Molecular Dynamics Simulation: The Langevin Piston Method. J. Chem. Phys. 1995, 103 (11), 4613–4621. https://doi.org/10.1063/1.470648.

(45) Andersen, H. C. Rattle: A “Velocity” Version of the Shake Algorithm for Molecular Dynamics Calculations. Journal of Computational Physics 1983, 52 (1), 24–34. https://doi.org/10.1016/0021-9991(83)90014-1.

(46) Lopez Cascales, J. J., Garro, A., Porasso, R. D., Enriz, R. D. The Dynamic Action Mechanism of Small Cationic Antimicrobial Peptides. Phys Chem Chem Phys 2014, 16 (39), 21694–21705. https://doi.org/10.1039/c4cp02537g.

(47) Alsop, R. J., Dhaliwal, A., Rheinstädter, M. C. Curcumin Protects Membranes through a Carpet or Insertion Model Depending on Hydration. Langmuir 2017, 33 (34), 8516–8524. https://doi.org/10.1021/acs.langmuir.7b01562.

(48) Marrink, S.-J., Berendsen, H. J. C. Simulation of Water Transport through a Lipid Membrane. J. Phys. Chem. 1994, 98 (15), 4155–4168. https://doi.org/10.1021/j100066a040.

(49) Hung, W.-C., Chen, F.-Y., Lee, C.-C., Sun, Y., Lee, M.-T., Huang, H. W. Membrane-Thinning Effect of Curcumin. Biophys. J. 2008, 94 (11), 4331–4338. https://doi.org/10.1529/biophysj.107.126888.

(50) Bennett, W. F. D., Tieleman, D. P. Computer Simulations of Lipid Membrane Domains. Biochimica et Biophysica Acta (BBA) – Biomembranes 2013, 1828 (8), 1765–1776. https://doi.org/10.1016/j.bbamem.2013.03.004.

(51) van Meer, G., Voelker, D. R., Feigenson, G. W. Membrane Lipids: Where They Are and How They Behave. Nat Rev Mol Cell Biol 2008, 9 (2), 112–124. https://doi.org/10.1038/nrm2330.

(52) Epand, R. M. Introduction to Membrane Lipids. In Methods in Membrane Lipids; Owen, D. M., Ed., Methods in Molecular Biology; Springer: New York, NY, 2015; pp 1–6. https://doi.org/10.1007/978-1-4939-1752-5_1.

(53) Baoukina, S., Tieleman, D. P. Computer Simulations of Phase Separation in Lipid Bilayers and Monolayers. Methods Mol. Biol. 2015, 1232, 307–322. https://doi.org/10.1007/978-1-4939-1752-5_21.

(54) Czolkos, I., Jesorka, A., Orwar, O. Molecular Phospholipid Films on Solid Supports. Soft Matter 2011, 7 (10), 4562–4576. https://doi.org/10.1039/C0SM01212B.

(55) Enkavi, G., Javanainen, M., Kulig, W., Róg, T., Vattulainen, I. Multiscale Simulations of Biological Membranes: The Challenge To Understand Biological Phenomena in a Living Substance. Chem. Rev. 2019, 119 (9), 5607–5774. https://doi.org/10.1021/acs.chemrev.8b00538.

(56) Sapay, N., Tieleman, D. P. Chapter 4 Molecular Dynamics Simulation of Lipid–Protein Interactions. In Current Topics in Membranes; Feller, S. E., Ed., Computational Modeling of Membrane Bilayers; Academic Press, 2008; Vol. 60, pp 111–130. https://doi.org/10.1016/S1063-5823(08)00004-5.

(57) Khajeh, A., Modarress, H. The Influence of Cholesterol on Interactions and Dynamics of Ibuprofen in a Lipid Bilayer. Biochimica et Biophysica Acta (BBA) – Biomembranes 2014, 1838 (10), 2431–2438. https://doi.org/10.1016/j.bbamem.2014.05.029.

(58) Pandit, S. A., Chiu, S.-W., Jakobsson, E., Grama, A., Scott, H. L. Cholesterol Surrogates: A Comparison of Cholesterol and 16:0 Ceramide in POPC Bilayers. Biophys J 2007, 92 (3), 920–927. https://doi.org/10.1529/biophysj.106.095034.

(59) Hub, J. S., Winkler, F. K., Merrick, M., de Groot, B. L. Potentials of Mean Force and Permeabilities for Carbon Dioxide, Ammonia, and Water Flux across a Rhesus Protein Channel and Lipid Membranes. J. Am. Chem. Soc. 2010, 132 (38), 13251–13263. https://doi.org/10.1021/ja102133x.

(60) Jedlovszky, P., Mezei, M. Effect of Cholesterol on the Properties of Phospholipid Membranes. 2. Free Energy Profile of Small Molecules. J. Phys. Chem. B 2015, 107 (22), 5322– 5332. https://doi.org/10.1021/jp021951x.

(61) Humphrey, W., Dalke, A., Schulten, K. VMD: Visual Molecular Dynamics. Journal of Molecular Graphics 1996, 14 (1), 33–38. https://doi.org/10.1016/0263-7855(96)00018-5.

(62) Bemporad, D., Essex, J. W., Luttmann, C. Permeation of Small Molecules through a Lipid Bilayer: A Computer Simulation Study. J. Phys. Chem. B 2004, 108 (15), 4875–4884. https://doi.org/10.1021/jp035260s.

(63) Issack, B. B., Peslherbe, G. H. Effects of Cholesterol on the Thermodynamics and Kinetics of Passive Transport of Water through Lipid Membranes. J Phys Chem B 2015, 119 (29), 9391–9400. https://doi.org/10.1021/jp510497r.

