## Supplementary Graphics for "A method for detection of permeation events in Molecular Dynamics simulations of lipid bilayers"

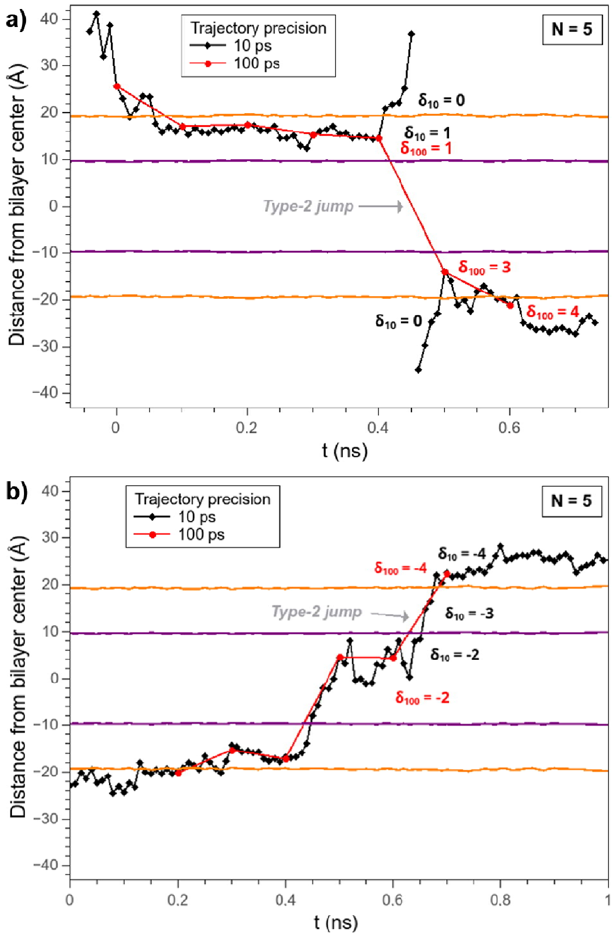


**Figure S1.** Influence of trajectory accuracy over permeation events detection. In both situations, the same trajectory is analyzed with precisions of 100 ps (red line) and 10 ps (black line). (a) Accepting the type-2 jump for a 100 ps precision trajectory leads to the register of a false positive event since the more detailed 10 ps record reveals instead the molecule left the bilayer, crossed the aqueous phase through the box edges, and entered the bilayer again by the opposite monolayer. (b) Although a type-2 jump to 100 ps precision occurs, the 10 ps detailed trajectory shows that the molecule indeed performs a permeation event.

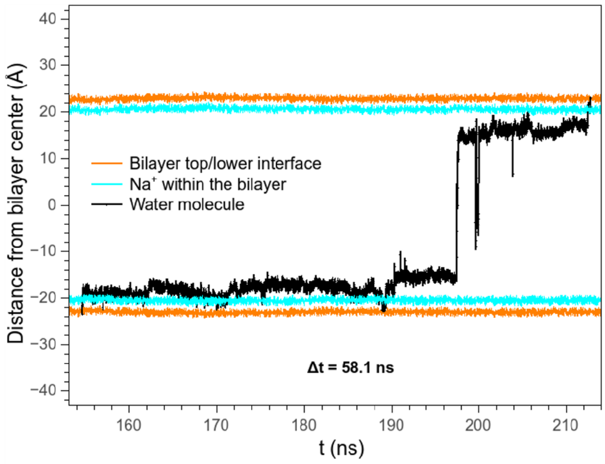


**Figure S2.** A Δt = 58.1 ns permeation event (black line) in the POPG bilayer (GG simulation set). The trajectory shows that the water molecule interacts for a long time with the Na+ ions inside the bilayer (cyan line). To calculate the average position of the ion plane at each instant, only the Na+ molecules within the bilayer (delimited by the orange lines) were considered.
